# A Portable and Scalable Genomic Analysis Pipeline for *Streptococcus pneumoniae* Surveillance: GPS Pipeline

**DOI:** 10.1101/2024.11.27.625679

**Authors:** Harry C. H. Hung, Narender Kumar, Victoria Dyster, Corin Yeats, Benjamin Metcalf, Yuan Li, Paulina A. Hawkins, Lesley McGee, Stephen D. Bentley, Stephanie W. Lo

## Abstract

Ever increasing global sequencing capacity provides an unprecedented opportunity in utilising genomic information captured from whole-genome sequencing to enhance pathogen surveillance. However, there is a growing need for developing user-friendly tools to effectively analyse the increasing volume of data. To meet this need, we have developed a genomic analysis pipeline, GPS Pipeline, which is portable and scalable to analyse genomes of *Streptococcus pneumoniae*, a major bacterial pathogen that is estimated to cause 317,000 child deaths worldwide every year. The GPS Pipeline is based on Nextflow and containerisation technology, and designed to enable researchers generating public health relevant output, including *in silico* serotypes, pneumococcal lineages (i.e. GPSCs), multilocus sequence types, and antimicrobial susceptibilities against 20 commonly used antibiotics,with minimal software setup requirements and bioinformatic expertise, in order to analyse genomic data at scale with ease. The GPS Pipeline provides a streamlined workflow that improves responsiveness in genomic surveillance on pneumococci.

**Data Summary:** The GPS Pipeline is available on GitHub at <u>github.com/GlobalPneumoSeq/gps-pipeline</u>. Published data from the GPS Database is available on Monocle Data Viewer at <u>data.monocle.sanger.ac.uk</u> and associated sequence read files are searchable and downloadable in the European Nucleotide Archive at <u>ebi.ac.uk/ena</u> via their ERR accession numbers.

**Impact Statement:** The GPS Pipeline advances global genomic surveillance of *Streptococcus pneumoniae* by providing a scalable, portable, and user-friendly tool for analysing whole-genome sequencing data. Leveraging Nextflow and containerisation technology, it minimises bioinformatics expertise requirements and infrastructure needs, making it particularly valuable in low- and middle-income countries where pneumococcal disease burden is high. This pipeline ensures reproducibility and stability across platforms, facilitating rapid and accurate pneumococci genomic analysis. By streamlining data processing, the GPS Pipeline enhances pathogen surveillance, generates evidence to support vaccine strategy development, and empowers researchers worldwide, ultimately contributing to improved public health outcomes.

## Introduction

Genomics has proven to be an effective tool to enhance pneumococcal disease surveillance and guide vaccine strategies as demonstrated by the Global Pneumococcal Sequencing (GPS) project [1]. With increasing accessibility to whole-genome sequencing (WGS), there is an unprecedented rise of publicly available pneumococcal genomes, as illustrated by the increasing number published on the European Nucleotide Archive (ENA) (Figure 1). This trend is likely to continue as it is reported that sequencing capacity has grown across the globe during the COVID-19 pandemic, and can now be more routinely applied on the surveillance of infectious diseases as we enter the post-COVID-19 period [2].

**Figure 1.**
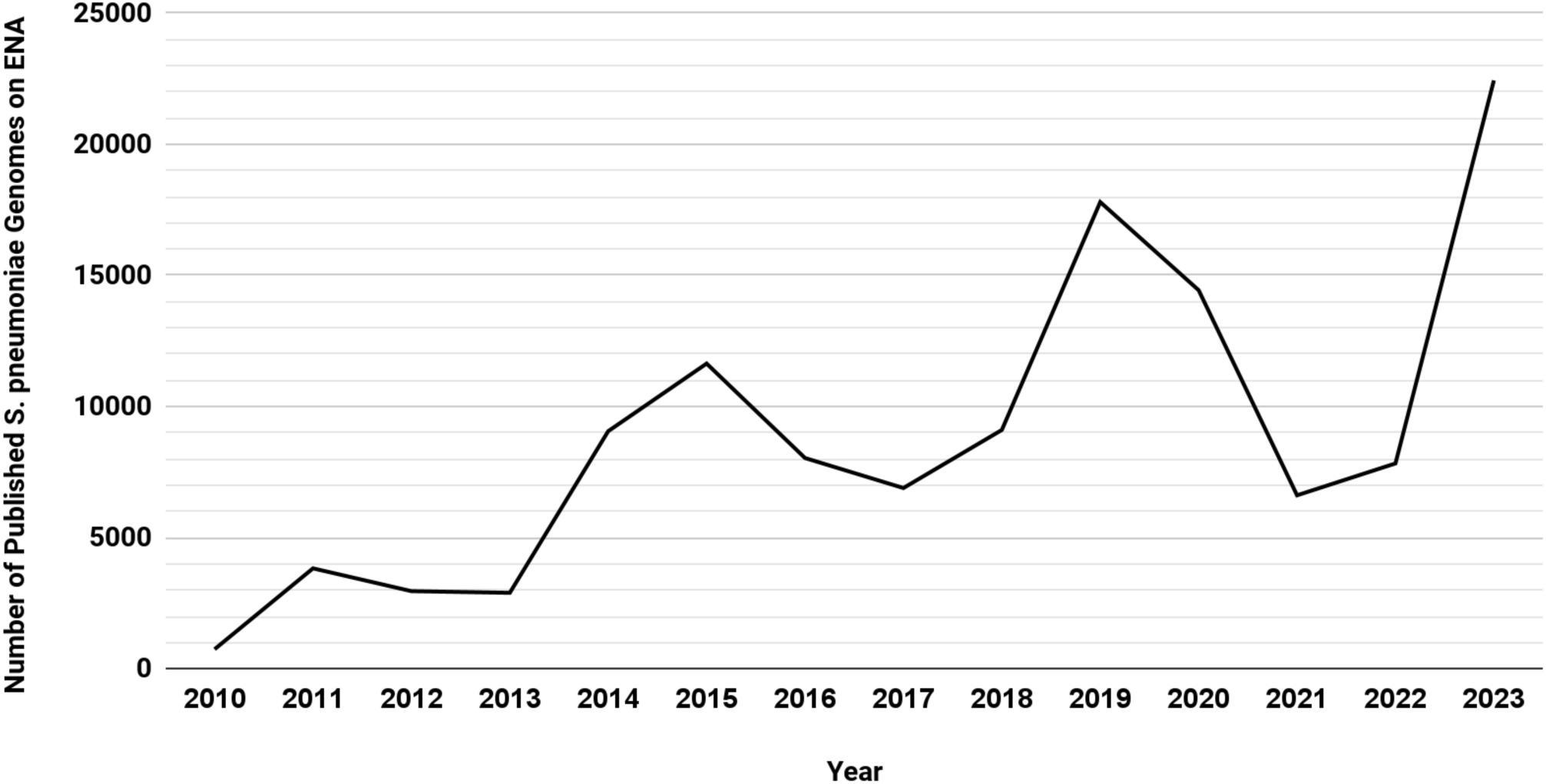
The number of *Streptococcus pneumoniae* samples available on ENA has been steadily rising over the years. Metadata of samples with NCBI Taxonomy ID of *Streptococcus pneumoniae* (1313) were retrieved from the ENA Browser on 9^th^ April 2024. Annual published genomes from Year 2010 to 2023 were computed based on the “first_public” field in the metadata. While the number of genomes does not consistently increase every year, there is an overall upward trend of publication of pneumococcal genomes.

Conducting genomic surveillance of pathogens requires an unique cross-disciplinary blend of knowledge in epidemiology, clinical microbiology, bioinformatics and IT infrastructure, in order to analyse the genomic data at scale and to understand the implication of the analysis results. However, there often exists a gap in bioinformatics expertise, especially in low- and middle-income countries (LMICs) [3]. At the same time, LMICs are the most affected by disease due to *Streptococcus pneumoniae* (pneumococcus), one of the leading causes of death in children under 5 years old [4].

While numerous projects, including the GPS project, have set out to assist the development of bioinformatics expertise and infrastructure in LMICs in the long term, it will take time for these efforts to come to fruition. One possible alternative approach to address these gaps could be the development of easy-to-use bioinformatics pipelines and tools. This approach would also benefit regions with more advanced bioinformatics capabilities by further enhancing their efficiency in processing genomic data. While well-known pipelines and tools exist, their constraints, such as difficulties in deployment and limited coverage of pneumococcus-specific analysis, hinder effective use for the pneumococcus surveillance in LMICs.

One of the first pipelines is the CDC pneumococcal typing pipeline developed by the *Streptococcus* Laboratory of CDC [5]. It is an automated Perl based pipeline that performs *in silico* typing and predicts antimicrobial susceptibilities. While it has been extensively used by CDC and some research groups to generate results for their publications, the deployment and usage of the pipeline are not clearly documented. The declining popularity of Perl, coupled with the environment-dependent nature of the pipeline, hinder its accessibility and usability to most researchers. In addition, the pipeline assumes that the input genome data has already undergone quality control. Consequently, users are responsible for performing quality control independently, a process that may require bioinformatics expertise.There is an effort to modernise the pipeline by refactoring it into a Nextflow pipeline, which promises to improve the ease of deployment, however it is still under development [6].

Another pipeline, Bactopia, is an all-encompassing bacterial genomic Nextflow pipeline with additional workflows called Bactopia Tools that can carry out a wide range of analysis with lots of flexibility [7]. However, due to its flexibility, it requires intricate configurations and preparations by researchers to instruct the pipeline to generate relevant information for pneumococcal surveillance. This characteristic lowers its readiness for quick deployment. Another limitation of Bactopia is its lack integration with PopPUNK, which is a tool widely used in the pneumococcus research community for pneumococcal lineage assignment (i.e. Global Pneumococcal Sequence Cluster, GPSC)[8].

Pathogenwatch [9], a web application designed for genomic surveillance, allows researchers to upload reads or assemblies of samples along with metadata to its website, where processing is performed on its servers. The results are then accessible and downloadable on the website. While it is a powerful and easy-to-use tool for genomic surveillance requiring minimal infrastructure, it requires a stable internet connection, which some LMICs may not have access to. As the data is processed remotely, researchers need to upload a large amount of data for each sample, which might not be feasible in areas that have slow and unreliable internet service. While Pathogenwatch provides detailed technical descriptions and open sourced the components, it is not designed to be deployed by users without web infrastructure expertise. It limits the ability of researchers to run their own instance locally.

To overcome the current challenges, we built an intuitive, portable, all-in-one pipeline to analyse pneumococcal genomes generating key public-health information including *in silico* serotypes, pneumococcal lineages (i.e. GPSC), multilocus sequence type (MLST), antimicrobial susceptibilities against 20 commonly used antibiotics, all at scale with a reasonable turnaround time.

## Theory and Implementation

To ensure portability, reproducibility, and ease of deployment, the GPS Pipeline (Figure 2 for schematic overview; Figure S1 for technical flowchart) was designed and implemented in Nextflow [10] within containerised environments deployed by Docker [11] as default or by Singularity [12] as an alternative option. The pipeline has a single entry point for inputting the directory containing raw reads of pneumococcal genomes (Illumina paired-end short read FASTQ files), then the FASTQ files are automatically processed by multiple Nextflow processes in parallel. The pipeline output includes 1) a single CSV file named results.csv containing all quality control (QC) and *in silico* typing results including *in silico* serotypes, GPSC, MLST, and antimicrobial susceptibilities to 20 common antibiotics in MIC value with Clinical & Laboratory Standards Institute (CLSI) guideline interpretation, 2) *de novo* assemblies in the format of FASTA files, and 3) a text file named info.txt that includes runtime information and software versions.

**Figure 2.**
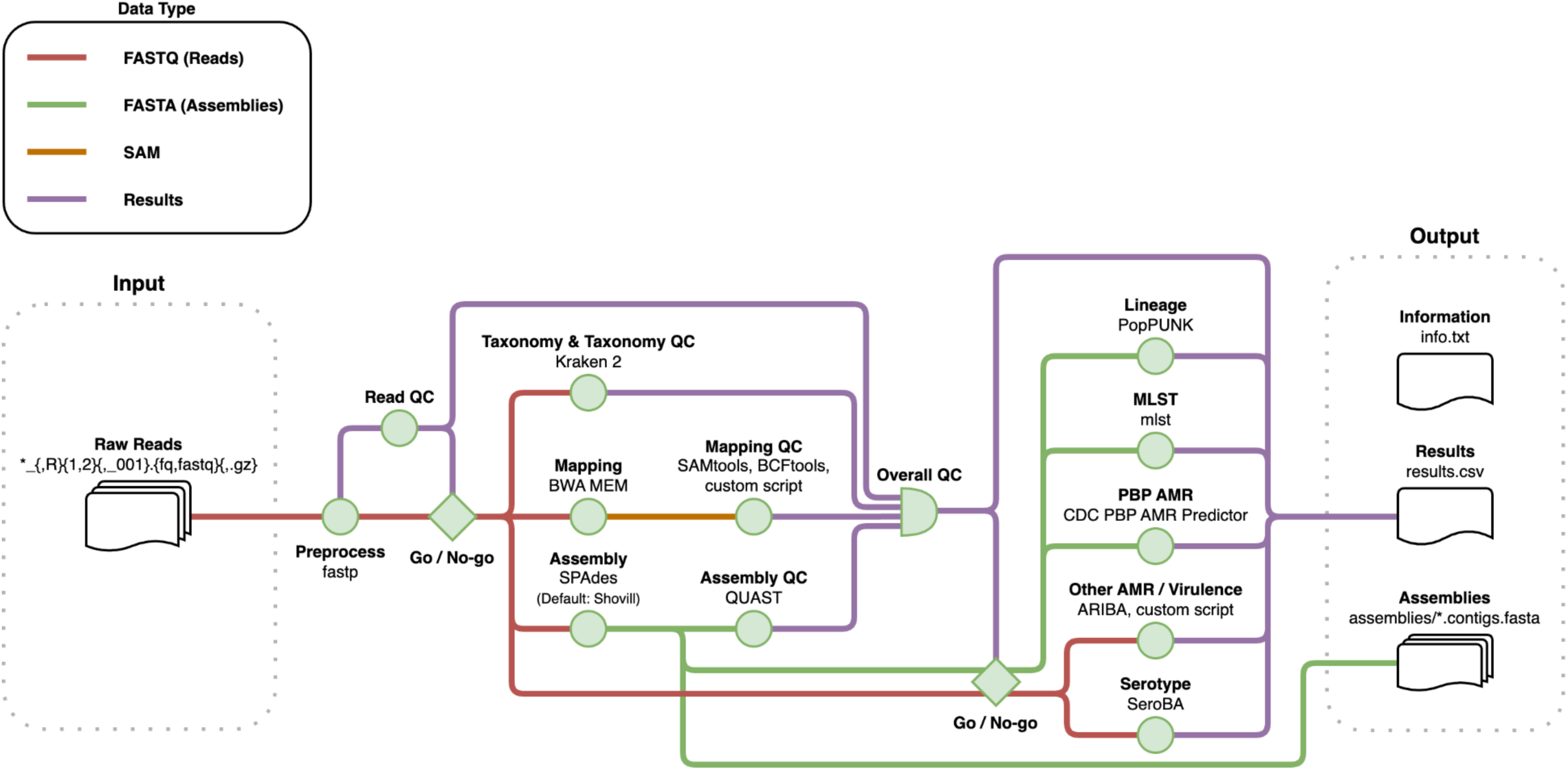
The schematic workflow of the GPS Pipeline. Once the user starts the GPS Pipeline with a directory path provided, the pipeline will capture all samples which raw read files match the glob pattern in that specific directory. All captured samples are processed by several bioinformatics tools and have their quality assessed by the quality control (QC) processes which jointly determine their overall quality based on the metrics of read, taxonomy, mapping and assembly. For samples that have passed the quality control, they are subjected to further analysis by the *in silico* typing processes which assign GPSC lineage, and predict serotype, MLST and a wide range of AMR and virulence targets. The results for all samples are collated into a single CSV file. The pipeline saves the runtime information and version of the software for reproducibility and the generated assembly of each sample.

The pipeline can be run without installation or administrative privileges on any POSIX-compatible system. It requires Bash 3.2 or above, Java 11 or above, and either Docker or Singularity installed. Compatible operating systems include Linux, Windows (through WSL), macOS and LSF-based high-performance computing (HPC) clusters. It is also compatible with the Seqera Platform (previously known as Nextflow Tower). We tested the GPS Pipeline on Linux (Ubuntu 22.04), Windows (Windows 11 with Ubuntu 22.04 through WSL2), and macOS (Sonoma 14.4) machines with 16GB or more RAM, as well as LSF-based HPCs (Wellcome Sanger Institute Farm5 and Farm22 HPCs), and generated consistent results. After downloading the pipeline and running its initialisation function which downloads all required additional files and container images, the pipeline can be used without an internet connection.

The pipeline logic of the GPS Pipeline is programmed in Nextflow Domain-Specific Language 2 (DSL2) and supplemented by custom Groovy functions. The pipeline logic invokes bioinformatics tools for analyses, and executes Shell or Python 3 scripts for data and metadata manipulations.

In this pipeline, we selected bioinformatics tools that are automated, reliable, fast, memory-efficient, and output results in a standardised text-based format that can be parsed programmatically. Based on these criteria, we have chosen the below bioinformatics tools and built custom scripts when necessary.

### Quality Control

The pipeline starts with a series of quality control checks on the FASTQ files and genomes based on the configurable criteria listed in Table S3. The pipeline first checks for corrupted FASTQ files (files that cannot be decompressed or contain incomplete data). Samples with any corrupted files are not further processed. An error message to indicate which file is corrupted is shown in the result file (results.csv).

Intact paired-end FASTQ files are then assessed by fastp v0.23.4 [13]. fastp automatically performs adapter trimming, quality filtering, and other operations to cleanup the FASTQ files for downstream processing, as well as acquiring the total base count of the read files. To pass the QC, the total base count should exceed the multiplication of minimum sequence depth and lower assembly length limit, which is 38 megabase pairs (Mbps). Pre-processed samples that pass QC are then processed in parallel through taxonomy, mapping and *de novo* assembly processes as stated below for further quality control based on these metrics.

To detect any potential contamination, the pipeline runs Kraken 2 v2.1.2 [14] on the FASTQ file pairs against the Minikraken v1 database [15] as default for taxonomy classification. Kraken 2 runs faster and requires less memory when compared to Kraken 1, and it uses a capped-size database available, enabling it to run on the usually limited system resources available on personal computers (PCs). In contrast, alternatives like GTDB-Tk, with its Genome Taxonomy Database (GTDB), have memory requirement exceeding more than most PCs can support (Table S2). A QC-passed genome should have ≥60% reads mapped to *S. pneumoniae* while ≤2% reads mapped to non-pneumococci species.

To detect a mixture of two or more pneumococcal isolates in a single sample, the FASTQ file pairs are mapped to the reference genome *S. pneumoniae* ATCC 700669 (NCBI accession no. FM211187) by default, the standard reference since the beginning of the GPS Project, using the BWA-MEM algorithm of BWA v0.7.17 [16]. Its successor, BWA-MEM2, was not used due to its much higher memory requirement rendering it unsuitable to run on PCs [17]. The output SAM files are converted into BAM files and sorted using SAMTools v1.16 [16], and the reference coverage percentage is then calculated. The sorted BAM files are used to call the single nucleotide polymorphisms (SNPs), which are saved into VCF files using BCFTools v1.16 [16]. Heterozygous SNP (het-SNP) sites are then counted by a custom Python script (<u>bin/het_snp_count.py</u>). This script filters out het-SNPs that are within 50-bp proximity to avoid overestimating the het-SNPs. In this step, a QC-passed genome is expected to have a coverage of the reference genome ≥60% and a count of het-SNP ≤220, which these thresholds were determined to be good metrics of the purity of isolates across ∼20,000 pneumococcal genomes in the GPS project.

To assess the assembled genome quality based on sequencing depth, length and number of contigs, *de novo* assembly is carried out using Shovill v1.1.0 [18] by default on the FASTQ file pairs. The quality metrics are summarised by QUAST v5.0.2 [19]. QC-passed genomes are expected to have a sequence depth of ≥20x, length of assembly between 1.9 - 2.3 Mbps, and number of contigs ≤500.

At the end of each QC check above, a Shell script (<u>bin/validate_file.sh</u> for FASTQ files integrity, <u>bin/get_read_qc.sh</u> for basic read quality, <u>bin/get_taxonomy_qc.sh</u> for taxonomy-based purity check, <u>bin/get_mapping_qc.sh</u> for mapping-based purity check, <u>bin/get_assembly_qc.sh</u> for assembly quality) assigns QC pass/fail category based on the QC metrics. Another Shell script, <u>bin/get_overall_qc.sh</u>, then assigns genomes that have passed all QC checks to the overall QC pass category. QC-failed genomes do not proceed downstream for *in silico* typing.

### *De novo* Assembly

Shovill v1.1.0 is set as the default assembler, with an alternative assembler Unicycler v0.5.0 [20] available in case Shovill fails to generate optimal assemblies for some samples. The selection was based on our benchmark of three popular *de novo* assemblers with automated optimisations, which two are SPAdes-based assemblers [21]: Shovill v1.1.0 and Unicycler v0.5.0, and one is Velvet-based assembler [22]: VelvetOptimiser v2.2.6 [23]. The benchmark tested the assemblers on reads from different Illumina short-read sequencing platforms (Table 1), and was carried out using the default parameters of the assemblers, except instructing them to use all available threads on the machine and discarding contigs with less than 500 base pairs (bps). The benchmark showed Shovill is the best assembler for the pipeline for FASTQ file pairs generated by different Illumina sequencing technologies. Shovill only takes 25.3% of runtime and 70.5% of maximum memory to generate assemblies of similar quality, as compared to Unicycler, based on number of contigs and N50 metrics, which is more than sufficient for the downstream tools in the pipeline. At the same time, VelvetOptimiser requires 172.7% of runtime compared to Shovill while generating worse quality assemblies and is a less stable software, making it unsuitable for the pipeline.

**Table 1.** Performance and quality comparison between *de novo* assemblers (* VelvetOptimiser repeatedly crashed while assembling one of the samples)

| Sequencing System | Assembler | Performance metrics<br>(mean ± 95% CI) | | Assembly quality metrics<br>(mean ± 95% CI) | | |
| --- | --- | --- | --- | --- | --- | --- |
| | | Run Time<br>(seconds) | Maximum memory<br>usage (MB) | No. of contigs<br>($\geq 500$ bps) | N50 | Assembly length<br>not in 1.9 - 2.3 Mbps |
| Illumina MiSeq<br>(n = 10) | Shovill | 255.47 ± 153.13 | 2997.71 ± 463.42 | 57.60 ± 21.13 | 97883.00 ± 29008.98 | 0 |
| | Unicycler | 689.11 ± 307.04 | 4284.82 ± 1389.25 | 55.10 ± 12.39 | 101738.10 ± 28805.49 | 0 |
| | VelvetOptimiser | 406.49 ± 152.41 | 1271.56 ± 684.86 | 185.50 ± 107.66 | 33612.40 ± 12354.01 | 1 |
| Illumina HiSeq<br>(n = 10) | Shovill | 176.12 ± 53.84 | 3542.72 ± 476.95 | 60.10 ± 17.73 | 76842.80 ± 16158.96 | 0 |
| | Unicycler | 723.20 ± 246.76 | 5566.50 ± 1267.15 | 60.50 ± 13.40 | 77030.80 ± 14694.38 | 0 |
| | VelvetOptimiser | 578.53 ± 176.60 | 6085.60 ± 1996.64 | 90.40 ± 31.81 | 38519.50 ± 17710.91 | 2 |
| Illumina NextSeq<br>(n = 10) | Shovill | 157.25 ± 31.35 | 3764.99 ± 534.04 | 46.10 ± 11.60 | 88366.10 ± 18882.47 | 0 |
| | Unicycler | 804.63 ± 203.17 | 5734.37 ± 1592.26 | 47.00 ± 11.79 | 133739.20 ± 77563.12 | 0 |
| | VelvetOptimiser | 838.78 ± 293.64 | 2315.92 ± 864.58 | 114.40 ± 50.92 | 18549.30 ± 11195.64 | 4 |
| Illumina NovaSeq<br>(n = 10; n = 9 for<br>VelvetOptimiser *) | Shovill | 117.60 ± 23.65 | 4219.44 ± 844.23 | 37.20 ± 7.70 | 173587.00 ± 83611.64 | 0 |
| | Unicycler | 573.42 ± 204.64 | 5022.16 ± 1597.11 | 38.90 ± 9.51 | 177481.30 ± 87313.33 | 0 |
| | VelvetOptimiser | 435.33 ± 184.84 | 3067.10 ± 1437.43 | 136.56 ± 106.57 | 45634.33 ± 12066.71 | 1 |

### In silico Typing

#### Genomic Lineage

We assign Global Pneumococcal Sequence Clusters (GPSCs) to define lineages which are based on genome-wide variations as compared to the conventional multilocus sequence typing (MLST) scheme which only takes the sequences of 7 housekeeping genes into account [8, 24]. GPSCs assignment is coherent by nature, allowing comparison of results between different studies. To assign GPSCs, PopPUNK v2.6.3 [25] and the latest PopPUNK GPS database v9 are used.

#### *In silico* Serotyping

For *in silico* serotype prediction, SeroBA v1.0.7 [26] was selected. It has a low memory and computational power requirement, yet high prediction accuracy and specificity. These advantages are evident when compared to PneumoCat which is another common *in silico* pneumococcal serotyping tool [27]. The current pipeline utilises SeroBA database v1.0.7. As new serotypes are detected, the pipeline can be easily updated to accommodate the latest version of SeroBA and its database.

#### MLST

mlst v2.23.0 [28] is used to infer MLST profile and sequence type (ST) from assemblies. The current pipeline uses pneumococcal PubMLST data as of 1st July 2024. As new ST are detected, the pipeline can also be updated to include the latest version of the PubMLST database.

#### Antimicrobial Resistance (AMR)

The GPS Pipeline predicts antimicrobial susceptibilities against 20 commonly used antibiotics using CDC PBP AMR Predictor [5] and ARIBA v2.14.6 [29], and interprets the MIC predictions based on the 2014 Clinical and Laboratory Standards Institute (CLSI) guideline [30]. The known resistance genes and mutations are summarised in Table 2.

**Table 2.**
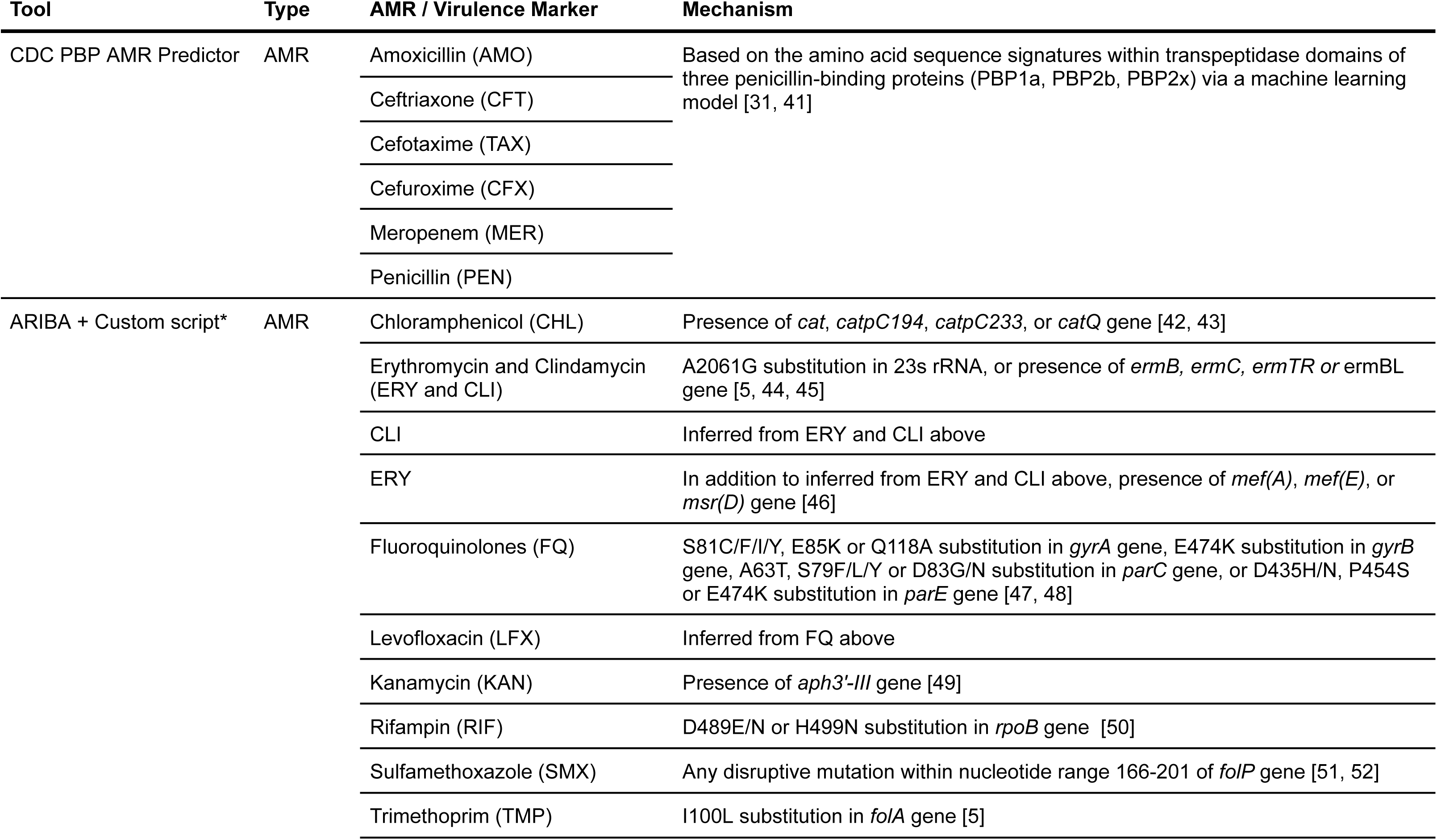
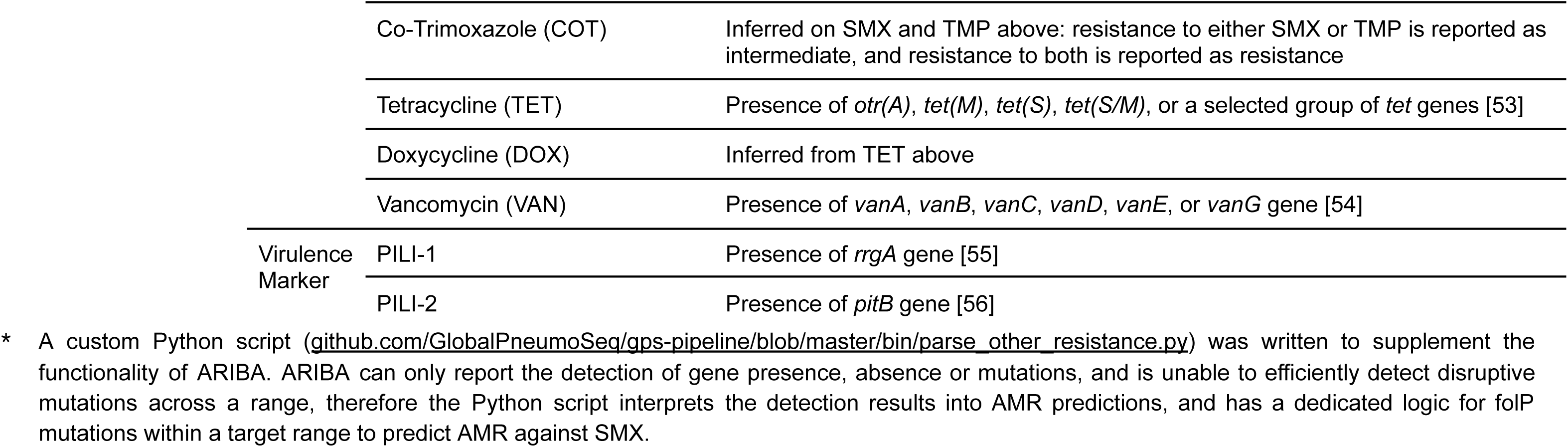
Tools and mechanisms deployed by the pipeline to predict antimicrobial resistances and virulence markers.

The CDC PBP AMR Predictor assigns PBP transpeptidase domain protein sequence types to the PBP1a, PBP2b, and PBP2x proteins encoded by draft genome assemblies, collectively referred to as PBP type. It then predicts the minimum inhibitory concentrations (MICs) against 6 β-lactam antibiotics, including: amoxicillin (AMO), ceftriaxone (CFT), cefotaxime (TAX), cefuroxime (CFX), meropenem (MER), and penicillin (PEN) using a machine learning model [5]. This machine learning model was previously shown to yield high percent essential agreement (MICs agree within ±1 dilution) >97% and percent category agreement (interpretive results agree) >93% in a pneumococcal genome dataset that was not used for building the machine learning model [31]. The resistance category against each antibiotic is also inferred from the predicted MIC according to the CLSI guidelines.

ARIBA v2.14.6 is used to perform the AMR predictions against 14 non-β-lactam antibiotics by detecting the presence or absence of known resistance genes or mutations in Table 2. It was chosen because newly recognised resistance genes or mutations can be readily added to the ARIBA database for detection, enabling the pipeline to quickly adapt to newly discovered AMR mechanisms in the future. We compiled a pneumococcus-specific ARIBA AMR database to detect the changes and mobile elements that determine resistance against 14 non-β-lactam antibiotics (Table 2) except for changes in *folP*. The nonsense mutations across the nucleotide range 166-201 (amino acid residue 56-67) of folP, which results in resistance to sulfamethoxazole (SMX) [44, 45], cannot be readily represented in an ARIBA database. Therefore, ARIBA is only used to detect and extract any mutations in the *folP* gene, and a Python script (<u>bin/parse_other_resistance.py</u>) is run subsequently to detect any disruptive mutation within the nucleotide range.

To avoid spurious detection, the presence of a resistance gene is confirmed by the Python script <u>bin/parse_other_resistance.py</u>, which determines if the gene meets the criteria of ≥80% coverage and ≥20x read depth. For a resistance mutation, a ≥10x sequence depth threshold is used for confirmation by this script. The categorical prediction of antimicrobial susceptibility (susceptible, intermediate, resistant) against chloramphenicol (CHL), clindamycin (CLI), co-trimoxazole (COT), doxycycline (DOX), erythromycin (ERY), ERY and CLI, fluoroquinolones (FQ), kanamycin (KAN), levofloxacin (LFX), rifampin (RIF), SMX, tetracycline (TET), trimethoprim (TMP), and vancomycin (VAN) is then saved to a report. The MIC range based on the categorical prediction is also added based on the CLSI guidelines [30].

#### Virulence Factor

The GPS Pipeline also detects the presence and absence of virulence factors: PILI-1, and PILI-2 using ARIBA v2.14.6 with the identical criteria of detecting resistance genes (≥80% coverage and ≥20x read depth).

### Containerisation

To implement the above functions in containerised environments, we included publicly available Docker images (Table S1) developed by a communal effort led by the State Public Health Bioinformatics Group (StaPH-B) [32] whenever possible. These images are well maintained and tested to be stable, appropriately versioned (which enables version pinning and simple upgrade), and open-source. During the development of the GPS Pipeline, we have also made contributions toward this effort by updating and adding testing stages to the Dockerfiles for the Docker images of PopPUNK and ARIBA, which were promptly reviewed and added to become part of their releases. In addition, we built Docker images of CDC PBP AMR Predictor and SeroBA, as up-to-date images were not available. To handle the extensive usage of command line utilities and Python scripts in the pipeline, Network-Multi Tool Docker image by WBITT [33] and Pandas Docker image by Alexander Mancevice [34] were chosen, respectively.

## Performance Benchmarking and Validation

The GPS Pipeline itself is less than 5MB in size. The default databases which are downloaded and uncompressed on the first run use about 19GB of disk space. The container images disk size is about 13GB with Docker or about 4.5GB with Singularity. Hence the essential files to run the pipeline occupy only 23.5 - 32GB of space in total. To benchmark the versatility and performance on various platforms, we processed 100 QC-passed genomes in the GPS database on a 16-core Ubuntu-based OpenStack instance running on Intel Xeon Gold 6226R processors, and 500 genomes (400 QC-passed and 100-QC failed) on the LSF-based Wellcome Sanger Institute Farm5 HPC running on a mixture of Intel Xeon Gold and AMD EPYC processors. It took 2 hours 48 minutes and 1 hour 40 minutes (depending on cluster availability), respectively.

We validated robustness and accuracy of the pipeline by running it on all published samples of the GPS database (n = 20,924, last accessed on 14^th^ June 2024) which only contain QC passed and *in silico* typed samples. In the GPS database, the genomes were previously assembled by a Perl pipeline written by the Wellcome Sanger Institute Pathogen Informatics Team [35], composed of Velvet version 1.2.10, VelvetOptimiser version 2.2.5 [23], SSPACE version 2.0 [36], GapFiller version v1.11 [37] and SMALT version 0.7.4 [38]. For *in silico* typing, mlst_check version 2.1.1630910 [39] was used to assigned MLST, PneumoCaT v1.2 [40] or SeroBA v1.0.0 [26] was used to infer serotypes for data generated before and after 15^th^ June 2018 respectively, CDC pneumococcal typing pipeline (which includes CDC PBP AMR Predictor) [5] was used to detect AMR with supplementary screening by ARIBA version 2.14.4 [29], and GPSC was assigned with PopPUNK version 1.1.5 with GPSC reference database v4 [25]. The *in silico* typing results of the GPS database have been verified to have high co-concordance against the phenotypic data [8].

All genomes from the published GPS database were successfully processed by the pipeline, with 98.0% (20,506/20,924) of the genomes considered to be QC passed. Although all samples passed the read QC, some failed in the assembly, mapping, or taxonomy QC, or a combination of these. The failures are primarily due to tightened QC parameters (Figure 3).

**Figure 3.**
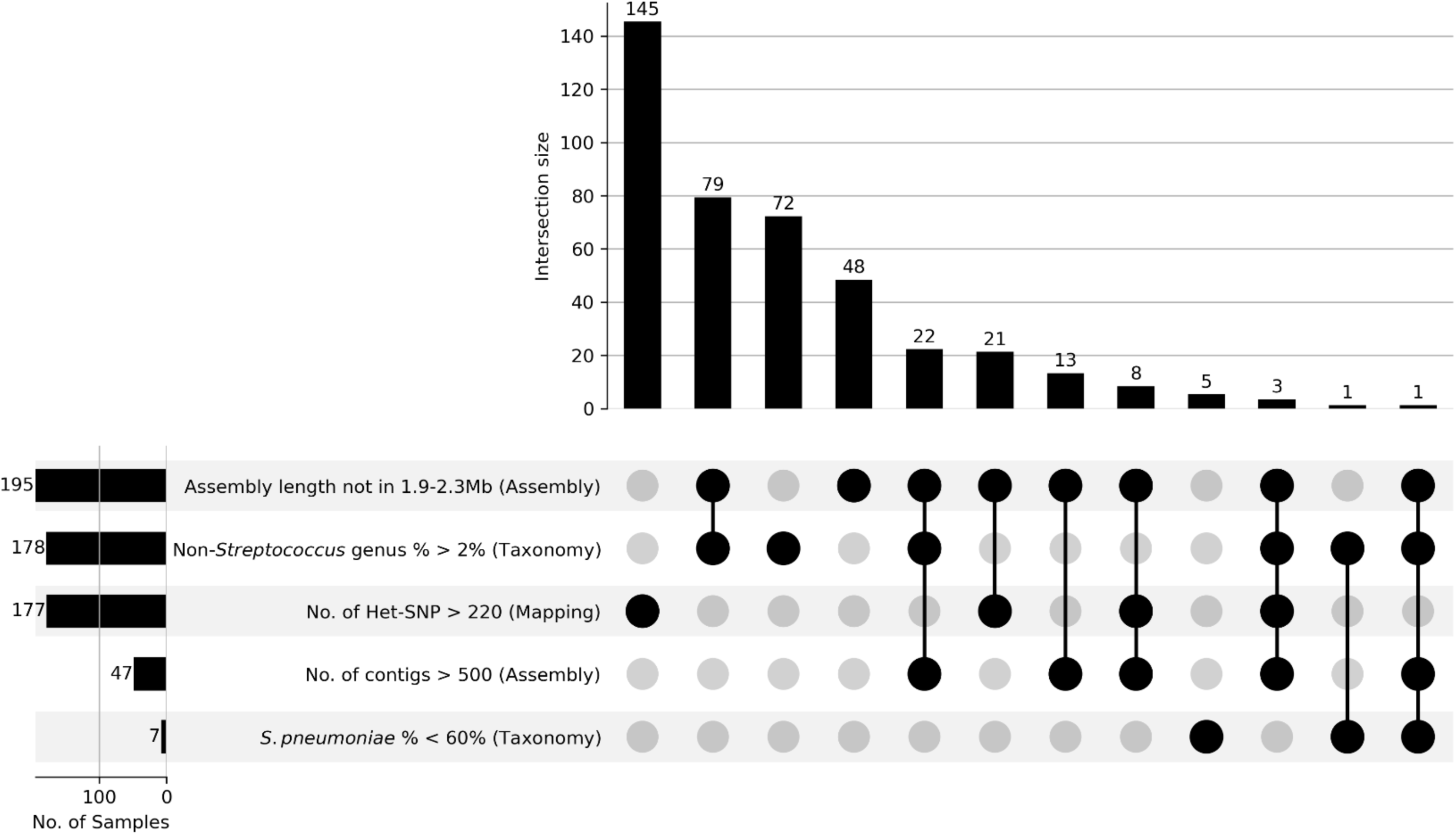
Reasons for 418 pneumococcal genomes failing QC by the GPS Pipeline. The UpSet plot shows the number of samples involved in each QC parameter failure in rows, and the number of samples in each combination of QC parameters failure in columns. A total of 418 samples failed in 604 QC parameters. Among failed samples, 83.7% (n = 350/418) failed in at least one of the new or updated QC parameters: non-*Streptococcus* genus percentage, and number of non-cluster heterozygous SNP (Het-SNP), suggesting the majority of failures are caused by tightened QC requirements.

The 20,506 QC-passed genomes were subject to *in silico* typing including 14 results shared by the GPS database and the GPS Pipeline: 1) GPSC; 2) MLST; 3) serotype; 4) PBP type and its inferred resistance category of 6 β-lactam antibiotics (AMO, CFT, TAX, CFX, MER, and PEN); 5-14) 10 individual predicted resistance category of non-β-lactam antibiotics (CHL, CLI, COT, DOX, ERY, ERY & CLI, FQ, LFX, RIF, and TET). Among all genomes typed by the GPS Pipeline, 92.6% (n = 19,002/20,506) have the identical results in the GPS database for all shared *in silico* typing results. For the remaining 1,504 genomes, they shared 1,710 changes in total (Table 3).

**Table 3.** Statistics of changes in 14 *in silico* typing categories in the GPS Pipeline when compared to the GPS Database.

| *in silico* Typing Categories | Description of Change | Count |
| --- | --- | --- |
| GPSC (n = 153) | Assignment of previously unassigned genomes | 146 |
|  | Failure to assign a previously assigned genomes | 6 |
|  | Updated assignment due to GPSC cluster merging event | 1 |
| MLST (n = 43) | Failure to assign a previously assigned genomes | 22 |
|  | Change in assignment | 13 |
|  | Assignment of previously unassigned genomes | 8 |
| Serotype (n = 280) | Change in assignment within same serogroup | 193 |
|  | Update of serotype name (i.e. 10X → 33G, 20 → 20A, 39X → 10D) | 39 |
|  | Failure to assign a previously assigned genomes | 34 |
|  | Change in assignment to a different serogroup | 14 |
| PBP type and its resulting resistance category of 6 β-lactam antibiotics (n = 74) | Change in PBP type; failure to assign previously resistance category | 30 |
|  | Change in PBP type; assignment of previously unassigned resistance category | 20 |
|  | No change in PBP type; change in resistance category | 11 |
|  | Change in PBP type; change in resistance category | 7 |
|  | Change in PBP type; no change in resistance category | 5 |
|  | No change in PBP type; failure to assign previously resistance category | 1 |
| CHL resistance category (n = 17) | Change from susceptible to resistant | 13 |
|  | Change from resistant to susceptible | 4 |
| CLI resistance category (n = 24) | Change from resistant to susceptible | 22 |
|  | Change from susceptible to resistant | 2 |
| COT resistance category (n = 159) | Change from susceptible to intermediate | 62 |
|  | Changed to indeterminable due to low gene coverage | 53 |
|  | Change from resistant to intermediate | 33 |
|  | Change from intermediate to susceptible | 6 |
|  | Change from intermediate to resistant | 5 |
| DOX resistance category (n = 38) | Change from resistant to susceptible | 38 |
| ERY resistance category (n = 215) | Change from resistant to susceptible | 190 |
|  | Change from susceptible to resistant | 25 |
| ERY & CLI resistance category (n = 24) | Change from resistant to susceptible | 22 |
|  | Change from susceptible to resistant | 2 |
| FQ resistance category (n = 191) | Change from intermediate to susceptible | 97 |
|  | Change from resistant to susceptible | 54 |
|  | Change from susceptible to intermediate | 37 |
|  | Changed to indeterminable due to low gene coverage | 2 |
|  | Change from susceptible to resistant | 1 |
| LFX resistance category (n = 366) | Change from susceptible to intermediate | 293 |
|  | Change from susceptible to resistant | 71 |
|  | Changed to indeterminable due to low gene coverage | 2 |
| RIF resistance category (n = 88) | Change from susceptible to resistant | 86 |
|  | Changed to indeterminable due to low gene coverage | 2 |
| TET resistance category (n = 38) | Change from resistant to susceptible | 38 |

For GPSC assignment, 0.75% of QC-passed genomes received a different result in the GPS Pipeline when compared to the GPS database. The majority is assignment of previously unassigned genomes as the latest PopPUNK GPS database used by the pipeline covers more lineages. The assignment failures can be attributed to changes in the PopPUNK algorithm between version 1.1.5 and 2.6.3, which is used to generate the GPS database and used by the GPS Pipeline, respectively.

For MLST assignment, the changes impact only 0.21% of genomes. They are a mixture of failure or change in assignments, and assignment of previously unassigned genomes. It is likely caused by the switch from mlst_check to mlst as the tool for MLST assignment, which has a different assignment algorithm.

For serotype prediction, 1.37% of genomes are predicted to have a different serotype. Most of them are changes within the same serogroup or updating to the latest naming conventions, it could either be the outcome of the change from PneumoCat to SeroBA, or improvements brought by an up-to-date serotype database in the latest SeroBA v1.0.7.

For PBP type assignment and β-lactam antibiotics resistance category prediction, 0.36% of genomes observe various changes. It is potentially due to the change of *de novo* assembler, as assemblies instead of sequence read files are passed to the CDC PBP AMR Predictor for typing and prediction, where changes in assemblies due to the assembler change could affect the prediction results.

For the non-β-lactam antibiotics resistance category prediction, 4.88% of genomes observed 1,160 changes in total. The changes are anticipated due to the move from CDC pneumococcal typing pipeline to a custom ARIBA-based prediction. As we have reimplemented the detection mechanism, updated the target genomic changes and mobile elements based on latest publications, and adjusted detection thresholds to avoid false positives, we are confident with the updated predictions.

## Discussion

A portable and adaptable pipeline for analysing *S. pneumoniae* genomes was built to extract key public health information including *in silico* serotype, GPSC, MLST and prediction of antimicrobial susceptibilities to 20 common antibiotics, with minimum requirement of IT infrastructure. The design of the pipeline allows users to analyse data ranging from high performance computing clusters (e.g. the GPS project at Sanger) to a laptop, yet retaining high reproducibility. The standardised *in silico* typing output from the GPS Pipeline could be easily adapted to be displayed on Microreact - an interactive web application to visualise epidemiological data and phylogeny, and well-suited for subsequent downstream processing and statistical analysis in R or other statistical software. The GPS core team and partners have used this pipeline to process >20,000 pneumococcal genomes since 2023, and the *in silico* typing output, combining the epidemiological metadata, are streamlined to upload to the project database, minimising the risk of manual errors. This demonstrates the GPS Pipeline’s capability to analyse data at scale, highlighting its potential for full automation, from processing raw sequencing data to generating comprehensive reports.

The GPS Pipeline was tested by over 15 research groups worldwide, issues and feedback collected over the past year were addressed and incorporated to develop this version. The pipeline has been generally well received, with some bug reports in the early phase of development which were promptly resolved as documented at the issues section in GitHub. We also received reports of difficulties initialising the pipeline in regions with unreliable internet service due to the large size of the databases. To address this challenge, we have worked with the developers of PopPUNK to generate a reduced-size PopPUNK database that brought the total database size from 30GB down to 19GB without a performance penalty. Most of the problems we received recently were related to setting up dependencies, particularly Java. We therefore added links to guides on how to set up the necessary dependencies of the pipeline to its manual in the GitHub repository (README.md).

Based on user feedback, once the pipeline is successfully set up, it stably processes the raw read files of their samples with ease. This includes users running the pipeline on their laptops, institute workstations, virtual machines on institute supercomputers, as well as HPCs. Therefore, it is reasonable to conclude that the GPS Pipeline has successfully accomplished its initial objective of becoming a portable and adaptable analysis pipeline for *S. pneumoniae* while remaining user friendly.

The future direction in pipeline development will explore the potential to accommodate read files from alternative sequencing technologies as the field expands beyond only using Illumina short-read sequencing technology.

## Conclusions

The GPS Pipeline allows scientists with limited bioinformatic skills to analyse their own pneumococcal genomes without the need for advanced computing resources. It readily extracts public-health relevant bacterial strain information to predict and evaluate the impact of pneumococcal conjugate vaccine impact on the pneumococcal population, thereby providing important information for the development of vaccine strategies. The automation of the pipeline enhances surveillance responsiveness. Its design, which is modularised and containerised, ensures portability, reproducibility, and simplifies future updates on bioinformatic tools. Additionally, it facilitates the expansion of databases for detecting new serotypes and antimicrobial resistance.

## Notes

https://github.com/GlobalPneumoSeq/gps-pipeline

https://data-viewer.monocle.sanger.ac.uk/project/gps

### Competing Interest Statement

The authors have declared no competing interest.

