## Supplemental Figure 1 for "A Portable and Scalable Genomic Analysis Pipeline for *Streptococcus pneumoniae* Surveillance: GPS Pipeline"

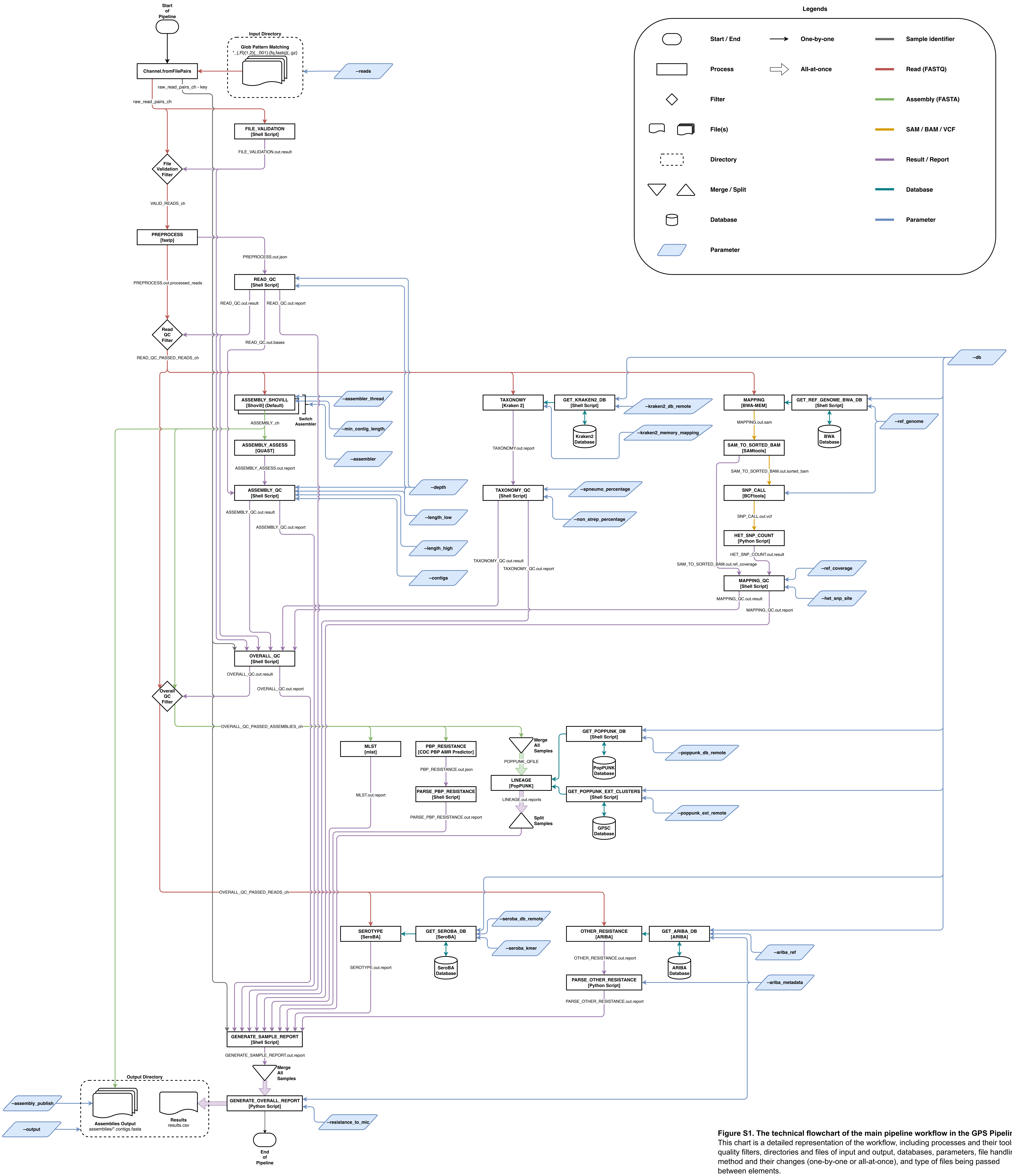


**Figure S1. The technical flowchart of the main pipeline workflow in the GPS Pipeline.** This chart is a detailed representation of the workflow, including processes and their tools, quality filters, directories and files of input and output, databases, parameters, file handling method and their changes (one-by-one or all-at-once), and type of files being passed between elements.
