## Supplemental Table 1 for "A Portable and Scalable Genomic Analysis Pipeline for *Streptococcus pneumoniae* Surveillance: GPS Pipeline"

**Table S1.** Details of Docker images used in the pipeline

| Docker Image Reference with Tag | Developer | Provided Tool(s) |
|---|---|---|
| amancevice/pandas:2.0.2 | Alexander Mancevice | Python 3.11.3; pandas 2.0.2 |
| sangerbentleygroup/seroba:1.0.7 | Bentley Group | SeroBA v1.0.7; SeroBA database v1.0.7 |
| sangerbentleygroup/spn-pbp-amr:23.10.2 | Bentley Group | CDC PBP AMR Predictor release 23.10.2 |
| staphb/ariba:2.14.6 | StaPH-B | ARIBA v2.14.6 |
| staphb/bcftools:1.16 | StaPH-B | BCFtools v1.16 |
| staphb/bwa:0.7.17 | StaPH-B | BWA v0.7.17 |
| staphb/fastp:0.23.4 | StaPH-B | fastp v0.23.4 |
| staphb/kraken2:2.1.2-no-db | StaPH-B | Kraken 2 v2.1.2 |
| staphb/mlst:2.23.0-2024-07-01 | StaPH-B | mlst v2.23.0; PubMLST database accessed on 1$^{st}$ July 2024 |
| staphb/poppunk:2.6.3 | StaPH-B | PopPUNK v2.6.3 |
| staphb/quast:5.0.2 | StaPH-B | QUAST v5.0.2 |
| staphb/samtools:1.16 | StaPH-B | SAMTools v1.16 |
| staphb/shovill:1.1.0-2022Dec | StaPH-B | Shovill v1.1.0 |
| staphb/unicycler:0.5.0 | StaPH-B | Unicycler v0.5.0 |
| wbitt/network-multitool:69aa4d5 | WBITT | BusyBox 1.34.1 with awk, bc, grep, gzip, paste, sed, sort, tar, zcat; GNU Bash 5.1.8; GNU Wget 1.21.2; jq 1.6 |
