## Supplemental Table 2 for "A Portable and Scalable Genomic Analysis Pipeline for *Streptococcus pneumoniae* Surveillance: GPS Pipeline"

**Table S2.** System requirements comparison between Kraken 2 and GTDB-Tk

| | **Disk Space** | **Memory** |
|---|---|---|
| Kraken 2 [57] | 8GB<br>(Minikraken v1 database) | 8GB<br>(Same size as the database) |
| GTDB-tk [58] | ~106GB<br>(GTDB) | ~90GB<br>(for Bacteria) |
