## Supplemental Table 3 for "A Portable and Scalable Genomic Analysis Pipeline for *Streptococcus pneumoniae* Surveillance: GPS Pipeline"

**Table S3.** Details of quality control parameters of the pipeline

| Type of Quality Control | Quality Control Parameter | Default Value |
|---|---|---|
| Read | Minimum base count<br>(not directly accessible, based on the multiplication of minimum assembly length and minimum sequencing depth) | ≥38,000,000 bp |
| Assembly | Maximum contig count | ≤500 contigs |
| | Minimum assembly length | ≥1,900,000 bp |
| | Maximum assembly length | ≤2,300,000 bp |
| | Minimum sequencing depth | ≥20x |
| Taxonomy | Minimum *S. pneumoniae* percentage in reads | ≥60% |
| | Maximum non-*Streptococcus* genus percentage in reads | ≤2% |
| Mapping | Minimum reference coverage percentage by the reads | ≥60% |
| | Maximum non-cluster heterozygous SNP (het-SNP) site count | ≤220 het-SNP sites |
